# Brachial Plexus Birth Injury Causes Location-Dependent Detriments in Glenohumeral Trabecular Bone Microstructure

**DOI:** 10.1101/2025.05.22.655580

**Authors:** Emily B. Fawcett, Nikhil N. Dixit, Katherine R. Saul, Jacqueline H. Cole

## Abstract

Brachial plexus birth injury (BPBI) is caused during a difficult childbirth when the head and neck are excessively stretched, and the brachial plexus nerve bundle is damaged. Injury causes lifelong arm impairment in 30-40% of those affected, but the extent to which bone microstructure is affected and how that relates to bone morphology alterations is unclear. Additionally, how injury location impacts trabecular bone, and the role limb disuse plays in these differential effects are unknown. Using two rat models of BPBI (postganglionic and preganglionic) and a disuse model (disarticulation), trabecular bone microstructure and glenohumeral morphology were characterized in the proximal humerus and distal scapula. For bone microstructure, the regions closest to the joint articulating surface incurred the greatest detriments with the least robust trabecular bone. These trabecular bone deficits in the humeral epiphysis and scapular neck were greater following preganglionic injury than postganglionic injury. The effects in the disarticulation group tended to align more with those in the postganglionic group, and greater detriments were observed in the scapula than in the humerus. Bone morphology metrics were explained mainly by quantity and architecture of trabecular bone but were not explained by the same metrics across all groups. Combining results with changes in bone morphology, data suggest differential effects with injury location and the effects of limb disuse to be most prominent in the postganglionic injury group and the distal scapula. In addition, the differences in results between postganglionic and preganglionic and macrostructural and microstructural deformity, suggest different drivers for postganglionic and preganglionic injuries and different factors causing macrostructural and microstructural alterations.

## 1. Introduction

Brachial plexus birth injury (BPBI) is one of the most common nerve injuries in children [1], caused by excessive stretching of the shoulder and neck during a difficult childbirth [2]. Injury results in damage to the brachial plexus nerve bundle and many co-morbidities, although injury mostly goes unrecognized by clinicians immediately following the birth [3]. Of the 1-3 in every 1,000 births in which injury occurs [1], 30 to 40 percent will sustain lifelong arm impairment [4]. Injury affects not only the way a child looks, with postural and joint deformities, but also results in limb disuse [5, 6], affecting how well the child is able to perform daily tasks such as eating and bathing. Damage to the nerve and the surrounding glenohumeral joint can be severe due to injury occurring during a period of rapid growth and development.

Clinically, BPBI portrays muscle weakness, contractures at the shoulder and elbow [7], joint dislocation [7], and an altered range of motion throughout the glenohumeral joint [8]. In animal studies, it has been shown that postural deformity is directly related to altered musculoskeletal loading of the joint [9]. However, only muscle changes have been investigated as potential mechanisms of altered loading, despite limb disuse being a known factor. Additionally, the severity of these injury characteristics changes with location of injury along the nerve [10]. When the nerve ruptures distal to the dorsal root ganglion (postganglionic injury), contractures and limb disuse are present [11]. When nerve avulsion occurs proximal to the dorsal root ganglion (preganglionic injury), the afferent signaling is preserved, and contractures at the shoulder and elbow are suppressed [12]. However, limb disuse still persists. Although injury location along the nerve portrays different abnormalities, prescribed treatments are typically the same and based on retrospective and case studies due to lack of knowledge around deformities [3]. At the extreme, surgeries to correct deformities have not been consistently successful [13].

Past studies have reported substantial morphological deformities at the glenohumeral joint following BPBI in rat models [9, 11, 14-16], due to altered mechanical loading of the joint from altered muscle growth [9]. In the scapula, bone alterations were more severe following postganglionic nerve injury, while in the humerus, they were more severe following preganglionic injury [17]. Comparing the two sites, the scapula underwent greater morphological changes than did the humerus [9, 17]. While the gross osseous deformities give insight into the shape of the bone post-injury, the trabecular microstructure, which has not been fully characterized, would allow insight into the strength of the bone.

Trabecular struts are essential in transferring loads from the joint surface to the cortical bone and model or remodel in response to altered mechanical loading [18], yet their reorganization following BPBI is not fully understood. A few studies have explored changes in trabecular microstructure within the proximal humeral head after a postganglionic nerve injury, showing that trabecular bone is less robust in the epiphysis [14, 15, 20]. We previously found that postganglionic injury also reduced trabecular bone volume in the scapular neck, with thinner and less connected trabeculae, and had a much greater effect there than in the humeral epiphysis. However, the effects of preganglionic injury on trabecular bone in the glenohumeral region, either in the scapula or the humerus, have not been examined in past studies.

The objectives of this study were to determine alterations in trabecular bone in the proximal humerus and lateral scapula due to altered loading following both postganglionic and preganglionic BPBI and to examine how these changes relate to macrostructural deformity and disuse of the affected limb. We hypothesized that trabecular bone loss would occur following BPBI, with postganglionic injury resulting in changes similar to disuse and worse than changes with preganglionic injury. Additionally, we hypothesized that trabecular bone changes would coincide with glenohumeral morphological changes. Investigation of trabecular microstructure throughout the glenohumeral region will provide a more complete picture of bone detriments that occur in response to altered mechanical loading following BPBI and reveal what trabecular bone metrics are most associated with the gross morphological changes seen at the glenohumeral joint.

## 2. Methods

### 2.1 Study Design

All animal surgeries and procedures were approved by the Institutional Animal Care and Use Committee at North Carolina State University. Male and female Sprague Dawley rat pups were split into four groups, including postganglionic neurectomy (n=22), preganglionic neurectomy (n=26), elbow disarticulation (n=12), and a sham control (n=15). Surgical interventions were performed under 2% isoflurane anesthesia at 3-5 days postnatally on one side (*affected*), with the contralateral side (*unaffected*) acting as an additional control. The postganglionic BPBI group underwent C5 and C6 nerve root excision distal to the dorsal root ganglion via incision of the pectoralis major, mirroring clinical nerve rupture [11]. The preganglionic BPBI group underwent C5 and C6 nerve root excision proximal to the dorsal root ganglion via supraclavicular incision, mirroring clinical nerve avulsion [6]. Due to surgical difficulty, five preganglionic neurectomies resulted in a partial nerve excision and were excluded from the study, leaving 21 subjects for the preganglionic group. As done in a previous study, partial nerve excisions were differentiated based on absence of the typical phenotype of an internally rotated shoulder and flexed wrist [6]. The disarticulation group underwent amputation at the elbow to mimic the limb disuse and gait patterns seen with BPBI but without nerve injury. The sham group underwent surgery similar to the postganglionic group, with incision via the pectoralis major muscle, but no nerve injury was induced. Incisions were closed with tissue adhesive and, for the disarticulation group, sutures.

During surgery, all groups were given a 2 mg/kg dose of bupivacaine injected subcutaneously in the shoulder region to serve as a localized nerve block. Immediately post-surgery, all groups were administered a 1 mg subcutaneous injection of saline and one prophylactic dose each of 0.02 mg/kg buprenorphine and 1.5 mg/kg carprofen injected subcutaneously. The disarticulation group received additional doses of meloxicam once daily for 5 days. Rats remained with their dam until weaning at postnatal day 21 and thereafter were housed up to 3 per cage. All animals were housed on a 12-hour light/12-hour dark cycle, were able to roam freely about their cages, and were given standard chow and water *ad libitum*. At 8 weeks, animals were euthanized via CO_2_ asphyxiation followed by bilateral thoracotomy. The torso and forelimbs were isolated using a rodent guillotine (Braintree Scientific, Inc., Braintree, MA), degloved, and then secured to a fixture to maintain a neutral shoulder posture and 90° of elbow flexion. The torso and forelimbs were fixed in 10% neutral buffered formalin for 48 hours, then immersed in 70% ethanol at 4°C for storage.

### 2.2 Micro-Computed Tomography

The bones from a subset of rats (sham: n=14; postganglionic: n=16, preganglionic: n=11, disarticulation: n=12) were scanned with micro-computed tomography (micro-CT) (µCT 80, SCANCO Medical AG, Brüttisellen, Switzerland). Bones were scanned in 70% ethanol at 70 kVp and 177 µA using a 0.5-mm Al filter, 800-ms integration time, 1,000 projections per rotation, and no frame averaging. Bone tissue density was analyzed using the SCANCO hydroxyapatite calibration phantom and setting a bone threshold of 250 mg/cm^3^ (2891 Hounsfield units). Three micro-CT scans were performed, one to analyze the overall bone morphology at the glenohumeral joint (macrostructure) and two to analyze the density and microstructure of trabecular bone in both the lateral scapula and proximal humerus. For macrostructure, the torso and forelimbs with intact glenohumeral joints were scanned and reconstructed at an isotropic voxel size of 36 µm. Each scan was thresholded for optimal appearance of bone surface details. For microstructure, humeri and scapulae were dissected and scanned separately and reconstructed at an isotropic voxel size of 10 µm, and these scans were reoriented so that the bones were aligned in the same anatomical position.

### 2.3 Microstructure Analysis

In the bone microstructure scans, volumes of interest (VOIs) were identified by manually contouring the trabecular bone in a series of slices and using automated morphing interpolation between slices in the SCANCO software. For the scapula, three VOIs were selected in secondary ossification centers in the glenoid fossa region near the joint articulating surface. This area was analyzed because of its rapid growth during the postnatal period [21] and because of the morphological changes previously seen along the articulating surface [9, 17]. The first VOI (*zone 1*) was identified in the subcoracoid secondary ossification center, starting at the articular surface and extending medially, parallel to the scapular spine axis, by 6.5% of the total scapular length, which was measured as the medial-lateral length along the scapular spine. The second VOI (*zone 2*) was located in the secondary ossification center of the upper glenoid, starting inferior to the subcoracoid ossification region and extending medially, parallel to the scapular spine axis, by 1.5% of the total scapular length. The third VOI (*zone 3*) was selected in the scapular neck, extending across the width of the neck and medially, parallel to the scapular spine axis, by 7.5% of the total scapular length. The VOI length percentages were determined in our previous study based on the largest length that could fit within each zone for the bone with the shortest scapular length; we confirmed that the same VOI length percentages would work for the scapular lengths in this study.

Two regions were selected for the humerus surrounding the proximal growth plate. This area was analyzed due to its proximity to the articulating surface. The epiphysis, the first humeral region, was identified directly superior to the proximal growth plate. It begins at the top of the humeral head and extends distally towards the growth plate by 12.5% of the total humeral length. The metaphysis, the second humeral region, was identified directly inferior to the proximal growth plate. It begins below the growth plate and extends distally down the bone by 5% of the total humeral length. Percentages were determined the same way in which the scapular zones were, with bones from a previous study.

Assessment of each scapular VOI and humeral region included analyses of the trabecular bone using standard metrics for bone quantity, property, and architecture [22]. These metrics included bone volume fraction (BV/TV), bone mineral density (BMD), tissue mineral density (TMD), connectivity density (Conn.D), trabecular number (Tb.N), trabecular thickness (Tb.Th), trabecular separation (TbSp), and degree of anisotropy (DA).

### 2.4 Macrostructure Analysis

Macrostructural measures were defined as gross morphological measurements taken from the lateral scapula and proximal humerus. Measurements included glenoid inclination angle (GIA), glenoid version angle (GVA), glenoid radius of curvature (GRC), humeral head thickness (HHT) and width (HHW), and humeral head radius of curvature (HRC), as described in a previous study [17]. Briefly, glenoid inclination angle analyzes the tilt of the scapula in the lateral-medial direction, with a smaller angle causing the scapula to be more medial. Glenoid version angle looks at the tilt of the scapula in the anterior-posterior direction with a smaller angle causing the scapula to tilt in the posterior direction. Curvature of the glenoid measured the radius of curvature within the articulating surface of the glenoid fossa and humeral head curvature measured the radius of curvature of the articulating surface along the humeral head [17]. Data for preganglionic were previously published in a study performed by collaborators [17].

Additionally, this study incorporated the scapular and humeral lengths for comparison between groups. Scapular length was measured as the distance from the glenoid fossa to the farthest medial point along the scapular spine axis. Humeral length was measured as the distance from the top of the humeral head to the most distal point.

### 2.5 Statistical Analysis

Normality of data was identified using D’Agostino-Pearson test. Limb comparisons, between the affected and unaffected limbs within a group, and group comparisons, between postganglionic, preganglionic, disarticulation, and sham were assessed. Trabecular bone metrics for limb comparisons were examined using paired parametric t-tests with Welch’s correction for unequal variances for data with a Gaussian distribution. A Wilcoxon matched-pairs signed rank test was performed for data without a Gaussian distribution. Group comparisons between affected-to-unaffected ratios were conducted via ANOVA with Tukey posthoc for those with data representative of a Gaussian distribution, and Kruskal-Wallis with Dunn posthoc for those with data not representative of a Gaussian distribution. The Tukey posthoc and Dunn posthoc tests acted to correct for comparison between multiple groups.

In addition, microstructural metrics from scapular zone 3 and the humeral epiphysis were used in a multiple regression analysis with macrostructural metrics from the same study. Microstructural metrics were assessed as predictor variables to explain previously gathered macrostructural metrics via stepwise multiple regression analysis with forward selection. The microstructural metrics were taken from the humeral epiphysis and scapular zone 3 (scapular neck) due to these regions lying along the articulating surface and portraying the least robust trabecular bone. All statistical analyses were performed with 95% confidence (α=0.05) in either Prism 6 (GraphPad Software, Inc., La Jolla, CA) or SAS 9.4 (SAS Institute Inc., Cary, NC).

## 3. Results

### 3.1 Effects on Bone Microstructure

#### 3.1.1 Limb Comparisons

Comparing the affected limb to the unaffected limb within groups, there were significant differences seen following postganglionic injury, preganglionic injury, and disarticulation at the lateral scapula and proximal humerus. Sham was generally unaffected throughout the bones. The scapula and humeral epiphysis portrayed similar differences between affected and unaffected limbs and the metaphysis seemed to show a lesser number of differences.

##### 3.1.1.1 Scapula

Alterations in trabecular microstructure of the affected limb occurred in all three zones of the scapula following postganglionic injury, preganglionic injury, and disarticulation, with zone 3 showing the greatest alterations (Figure 1, Table 1). The affected limbs in postganglionic, preganglionic, and disarticulation groups showed 13-16%, 11-29%, and 15-25% lower bone volume fraction respectively and 12-15%, 11-28%, and 16-24% lower bone mineral density respectively, in all zones. For tissue mineral density, postganglionic showed a 1-3% decrease on the affected side for zones 1 and 2, preganglionic showed a 2% decrease on the affected side for zones 2 and 3, and disarticulation showed a 2% decrease on the affected side for zone 2. Metrics for trabecular architecture were altered on the affected side in all three groups, but varied slightly between zones. In zone 1, postganglionic portrayed 8% lower trabecular thickness and 18% increased trabecular separation and preganglionic had 19% lower connectivity density, 17% lower trabecular number, 7% lower degree of anisotropy, and 27% increased trabecular separation. Disarticulation showed 17% lower connectivity density, 15% lower trabecular number, and 25% increased trabecular separation, and tended to show 17% decreased trabecular thickness (p=0.055), and 7% higher degree of anisotropy (p=0.056). Sham also had a slight change on the affected side, showing a trend towards a 3% decrease in trabecular thickness (p=0.082) in the affected limb. In zone 2, postganglionic portrayed 22% increased trabecular separation, and tended to show 7% lower trabecular thickness (p=0.075) in the affected limb (Figure). Preganglionic had 10% lower trabecular thickness and 11% lower degree of anisotropy, and 24% increased connectivity density and 6% increased trabecular number on the affected side. Disarticulation showed 15% decreased trabecular number and 12% decreased trabecular thickness, 25% increased trabecular separation, and tended to show 5% decreased degree of anisotropy (p=0.043) on the affected side. Sham did not have any differences.

**Table 1.** Percent differences in trabecular bone metrics between affected and unaffected limbs in scapular volumes of interest (VOI). **p < 0.05, *p < 0.1.

| Intervention | VOI | Affected Side versus Unaffected Side (%) |  |  |  |  |  |  |  |
| --- | --- | --- | --- | --- | --- | --- | --- | --- | --- |
|  |  | BV/TV | BMD | TMD | Conn.D | Tb.N | Tb.Th | Tb.Sp | DA |
| Sham | Zone 1 | -4.2 | -2.3 | 0.4 | 2.2 | -1.1 | *-2.5 | 3.1 | 1.6 |
|  | Zone 2 | 3.0 | 2.4 | 0.1 | 10.0 | 0.7 | 2.4 | -2.4 | 0.9 |
|  | Zone 3 | 1.5 | 1.6 | <u>0.9</u> | 1.4 | -1.4 | **2.3 | 2.9 | -0.2 |
| Postganglionic | Zone 1 | ** -13.3 | ** -12.4 | ** -1.1 | -4.4 | -8.5 | ** -7.8 | **18.0 | 4.3 |
|  | Zone 2 | ** -15.8 | ** -12.8 | ** -2.9 | -3.0 | -3.9 | * -6.5 | **21.9 | -2.9 |
|  | Zone 3 | ** -15.2 | ** -15.3 | -0.7 | ** -13.6 | ** -16.0 | 0.0 | **30.1 | ** -7.2 |
| Preganglionic | Zone 1 | ** -18.4 | ** -16.6 | -0.3 | ** -18.6 | ** -17.2 | -1.3 | **26.6 | ** -7.2 |
|  | Zone 2 | ** -11.4 | ** -11.1 | ** -2.0 | **24.4 | **6.4 | ** -9.8 | 7.1 | ** -11.1 |
|  | Zone 3 | ** -28.7 | ** -28.2 | ** -1.9 | ** -24.9 | ** -21.4 | ** -9.7 | **40.2 | -4.7 |
| Disarticulation | Zone 1 | ** -25.4 | ** -23.8 | 1.1 | ** -17.3 | ** -15.4 | * -16.6 | **24.7 | *7.2 |
|  | Zone 2 | ** -23.1 | ** -20.8 | ** -2.5 | 11.0 | ** -15.4 | ** -12.4 | **24.6 | * -4.5 |
|  | Zone 3 | ** -15.4 | ** -16.2 | 0.0 | ** -23.3 | ** -16.2 | **7.1 | **37.4 | ** -11.3 |

\*\*p < 0.05, \*p < 0.1.

**Figure 1.**
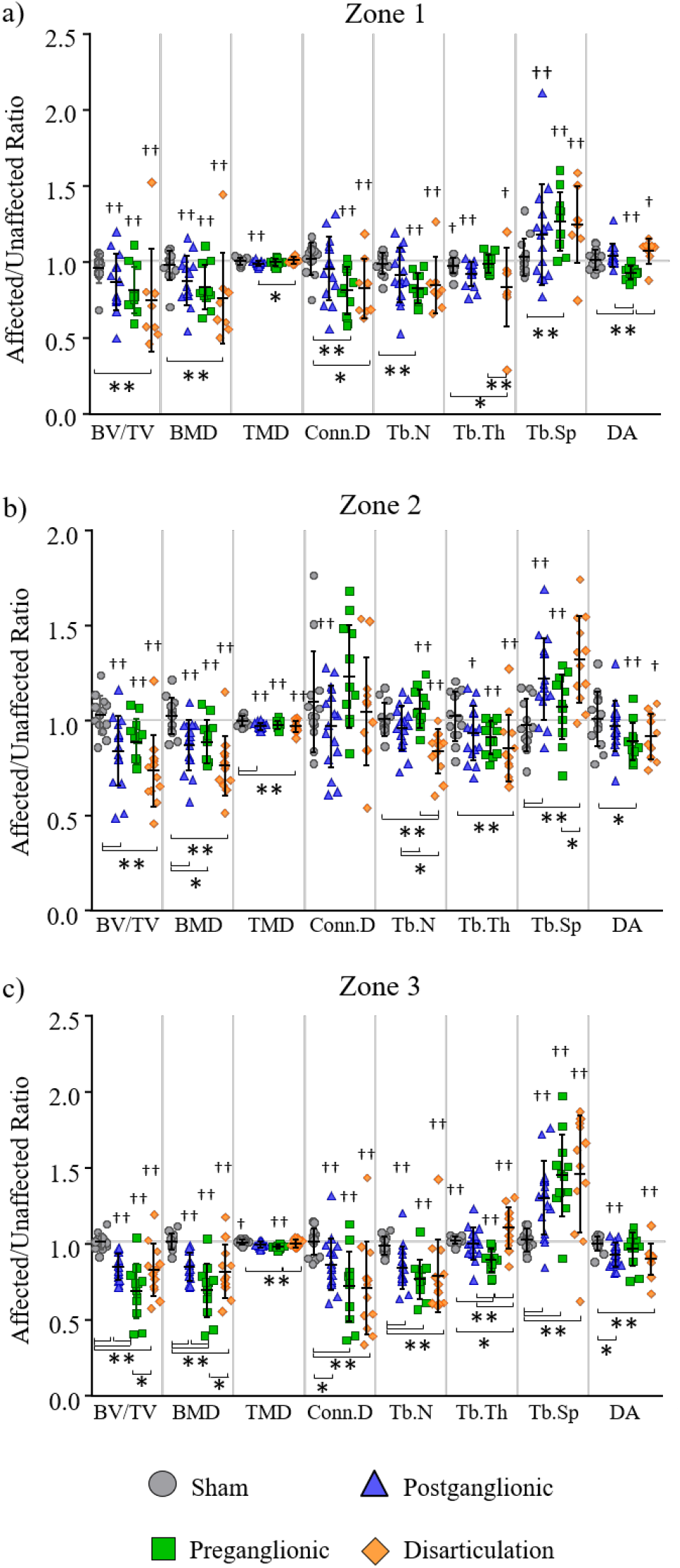
Trabecular bone metrics in the distal scapula. Altered trabecular microstructure was exhibited for zone 1 (a), zone 2 (b), and zone 3 (c) following postganglionic BPBI, preganglionic BPBI, and disarticulation, with zone 3 showing the most differences in all intervention groups. ^‡^p < 0.05, ^†^p < 0.10 affected vs. unaffected limb; **p < 0.05, *p < 0.10 groups differences.

Scapular zone 3 had the greatest number of differences between affected and unaffected limbs (Figure 1c). Bone quantity was again reduced in the three intervention groups (postganglionic, preganglionic, and disarticulation) with lower bone volume fraction (−15%, -29%, -15% respectively) and bone mineral density (−15%, -28%, -16% respectively). Material property was reduced by 2% in the affected limb in preganglionic and tended to be increased by 1% in the affected limb of sham (p=0.075). For trabecular architecture, postganglionic portrayed 14% reduced connectivity density, 16% lower trabecular number, and 7% decreased degree of anisotropy and 30% increased trabecular separation. Preganglionic had 25% lower connectivity density, 21% lower trabecular number and 10% decreased thickness, and 40% increased trabecular separation. Disarticulation was similar with 23% decreased connectivity density, 16% lower trabecular number, and 11% decreased degree of anisotropy and 7% increased trabecular thickness and 37% increase separation. Sham also showed a 2% increase in trabecular thickness on the affected side.

##### 3.1.1.2 Humerus

Trabecular microstructure alterations were present in all regions of the affected arm following postganglionic and preganglionic injury but was worse in the epiphysis (Figure 2, Table 2). Disarticulation presented with alteration in the epiphysis of the affected arm but did not show significant changes in the metaphysis. In the epiphysis ad metaphysis regions, postganglionic and preganglionic had 11-16% and 22-39% less BV/TV respectively and 8-16% lower and 18-40% lower BMD on the affected side respectively. TMD was decreased on the affected side by 1% and 2% in the epiphysis for postganglionic and preganglionic groups respectively. The disarticulation group showed a 7% decrease in BV/TV and BMD on the affected side in only the epiphysis. For trabecular architecture, many changes were seen in the epiphysis of the affected limb. Postganglionic and preganglionic resulted in 14% and 29% lower Conn.D, 11% and 28% lower Tb.N, 6% and 15% lower Thb.Th, and 14% and 52% increased Tb.Sp. Disarticulation resulted in decreased Tb.N by 5% and increased Tb.Sp and DA by 7% and 3% respectivley. Sham showed no differences.

**Table 2.**
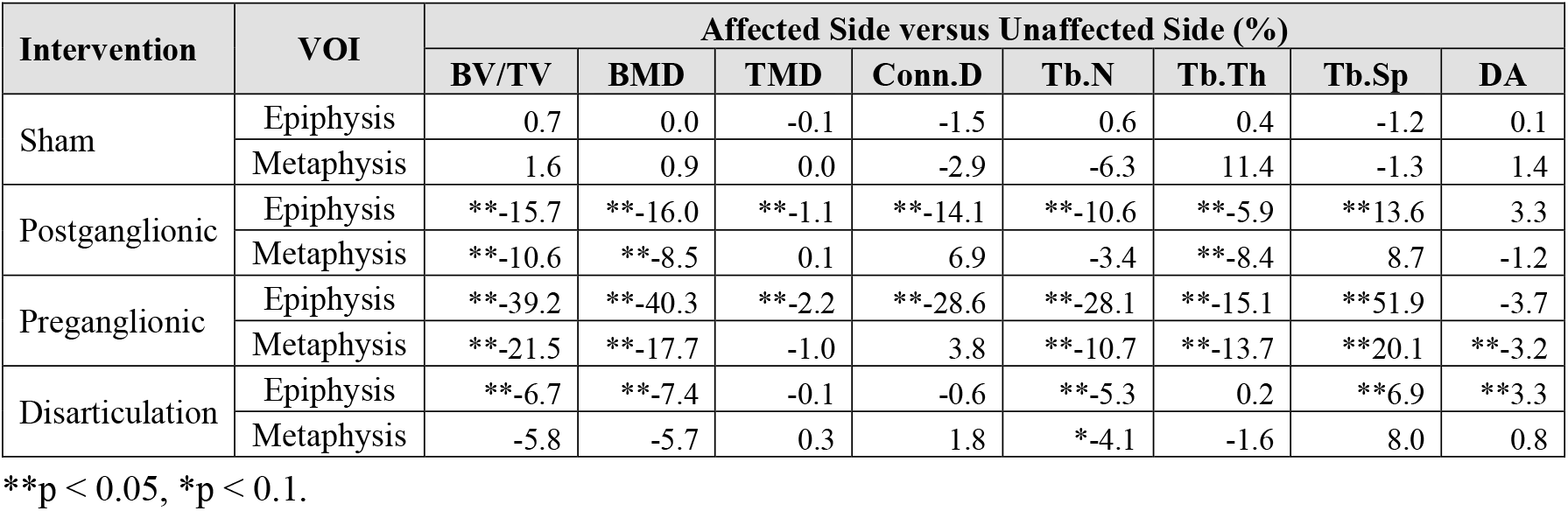
Percent differences in trabecular bone metrics between affected and unaffected limbs in humeral volumes of interest (VOI).

**Figure 2.**
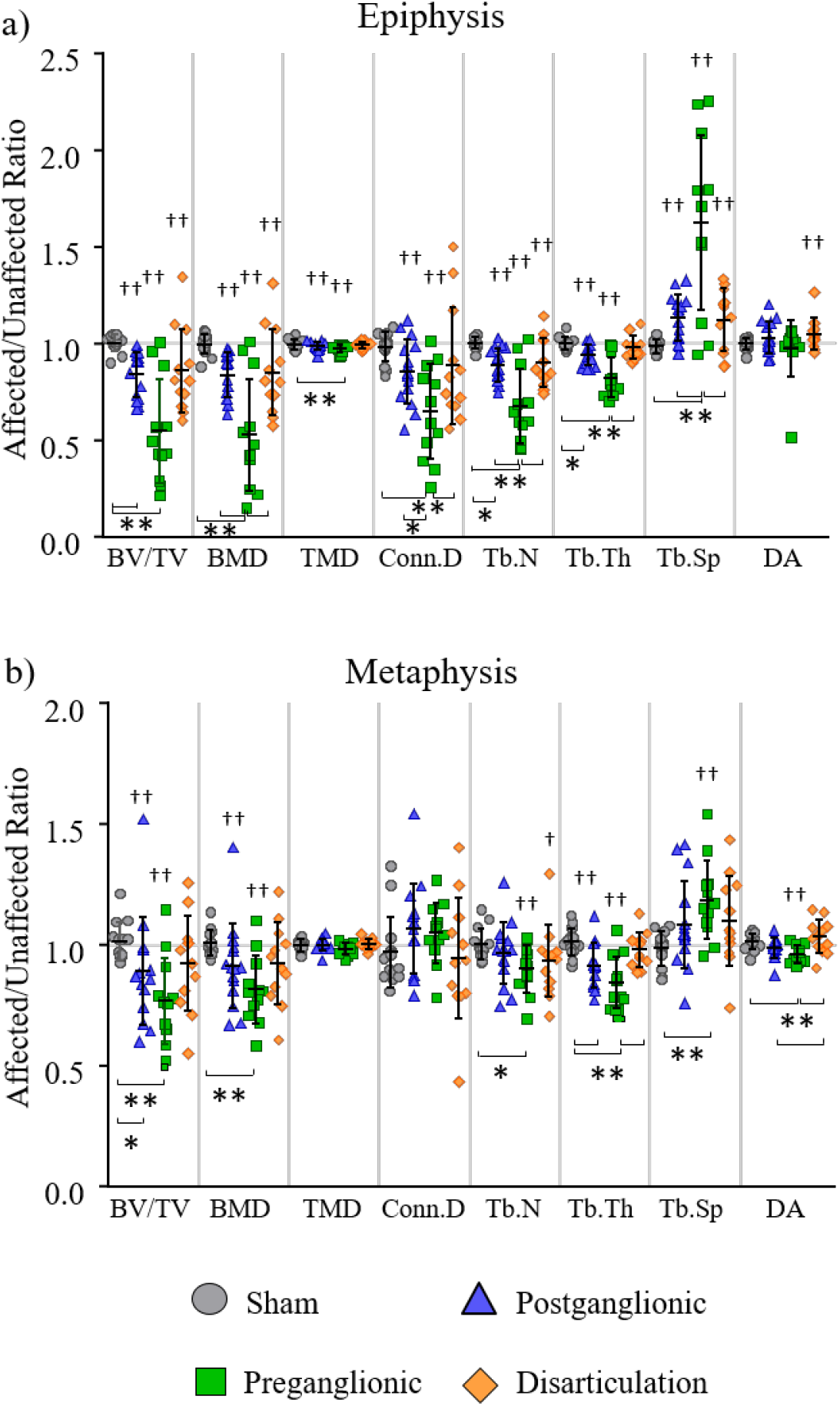
Trabecular bone metrics in the proximal humerus. Altered trabecular microstructure was exhibited for the epiphysis (a) and metaphysis (b) following postganglionic BPBI, preganglionic BPBI, and disarticulation with the epiphysis showing the most differences in all intervention groups. ^‡^p < 0.05, ^†^p < 0.10 affected vs. unaffected limb; **p < 0.05, *p < 0.10 group differences.

#### 3.2.2 Group Comparisons

Postganglionic, preganglionic, and disarticulation groups resulted in less robust trabecular bone compared to sham (Figure 3). Throughout both regions, postganglionic was most similar to sham.

##### 3.1.2.1 Scapula

Scapular zone 3 incurred the greatest changes in postganglionic, preganglionic, and disarticulation when compared to sham (Figure 1). In general, preganglionic exhibited worse trabecular metrics than postganglionic. Preganglionic and disarticulation were similar in zone 1. Zone 2 showed worse robustness in disarticulation and zone 3 showed the least robustness in preganglionic.

In zone 1, no postganglionic metrics showed differences compared to sham (Figure 1a). However, postganglionic tended to have less TMD (p=0.084, -2%) compared to disarticulation. Preganglionic showed lower Conn.D (−20%), Tb,N (−16%), and DA (−9%) and increased Tb.Sp (+23%) compared to sham. It also had significantly lower DA compared to both postganglionic (−11%), and disarticulation (−13%). Disarticulation showed worse BV/TV (−22%) and BMD (−22%) compared to sham and worse Tb.Th (−18%) compared to preganglionic. Disarticulation also tended to show less robust Conn.D (p=0.077, -19%) and Tb.Th (p=0.061, -14%) compared to sham.

Scapular zone 2 portrayed more differences than in zone 1 (Figure 1b). Compared to sham, postganglionic had lower BV/TV (−18%) and TMD (−3%), and increased Tb.Sp (+25%). Postganglionic also tended to have lower BMD (p=0.056, -15%) compared to sham and lower Tb.N (p=0.080, -10%) compared to preganglionic. For preganglionic, there were no significant differences compared to sham, but preganglionic tended to have lower BMD (p=0.010, -13%) and DA (p=0.096, -12%) compared to sham. Disarticulation showed lower BV/TV (−29%), BMD (−26%), TMD (−3%), Tb.N (−17%), and Tb.Th (−16%), and increased Tb.Sp (+35%) when compared to sham. Disarticulation also showed decreased Tb.N (−26%) compared to preganglionic, and tended to show lower Tb.N (p=0.020, -16%) and increased Tb.Sp (p=0.014, +19%) compared to postganglionic and preganglionic respectively.

The worst scapular region was zone 3, which lies nearest the articulating surface (Figure 1c). Postganglionic portrayed lower BV/TV (−16%), BMD (−17%), Tb.N (−15%), and Tb.Th (−2%), and increased Tb.Sp (+27%) compared to sham. It also tended to show less Conn.D (p=0.088, - 15%) and DA (p=0.059, -7%) compared to sham. Preganglionic had decreased BV/TV (−32%), BMD (−32%), TMD (−3%), Conn.D (−29%), Tb.N (−22%), and Tb.Th (−13%), and increased Tb.Sp (+42%) when compared to sham. In comparison to postganglionic, preganglionic had lower BV/TV (−19%), BMD (−18%), and Tb.Th (−11%). Compared to disarticulation, preganglionic had lower TMD (−2%) and Tb.Th (−19%), and tended to have lower BV/TV (p=0.052, -17%) and BMD (p=0.095, -15%). Lastly, disarticulation showed decreased BV/TV (−19%), BMD (−20%), Conn.D (−30%), Tb.N (−20%), and DA (−10%), increased Tb.Sp (43%), and tended to show increased Tb.Th (p=0.064, 8%) compared to sham.

##### 3.1.2.2 Humerus

The epiphysis portrayed greater differences between groups than the metaphysis. Preganglionic was the least robust of the groups and postganglionic and disarticulation were of similar robustness (Figure 2).

In the epiphysis, postganglionic exhibited lower BV/TV (−16%) and tended to have lower Tb.N (p=0.069, -11%) and Tb.Th (p=0.042, -6%) compared to sham (Figure 2a). Preganglionic showed lower BV/TV (−45%), BMD (−47%), TMD (−2%), Conn.D (−34%), Tb.N (−32%), and Tb.Th (−18%), and increased Tb.Sp (+65%) compared to sham. When compared to postganglionic, preganglionic portrayed less BMD (−37%) and Tb.N (−24%), increased Tb.Sp (65%), and tended to portray less Conn.D (p=0.059, -24%). Compared to disarticulation, preganglionic showed lower BV/TV (−38%), Conn.D (−27%), Tb.N (−25%), and Tb.Th (−16%), and increased Tb.Sp (+45%). The disarticulation group showed no significant differences between either sham or postganglionic.

Fewer metrics were affected in the metaphysis with mainly the preganglionic group showing detriments (Figure 2b). In postganglionic, there was lower Tb.Th (−10%) and a trend towards lower BV/TV (p=0.098, -12%) compared to sham, along with a trend towards decreased DA (p=0.054, -5%) compared to disarticulation. Preganglionic portrayed decreased BV/TV (−24%), BMD (−19%), Tb.Th (−17%), and DA (−5%), increased Tb.Sp (+20%), and tended to portray decreased Tb.N (p=0.051, -10%) compared to sham. Compared to disarticulation, preganglionic showed less Tb.Th (−14%) and DA (−7%). The disarticulation group had no significant differences when compared to the sham group.

### 3.2 Effects on Bone Macrostructure

Looking at the affected limbs within each group, postganglionic mostly showed deformity in the scapula, preganglionic mainly showed deformity in the humerus, and disarticulation showed some deformity in both the scapula and humerus (Table 3). The affected side in postganglionic portrayed a smaller GIA (−10 degrees), shorter humeral length (−2%), and tended to portray flattened GRC (p=0.064, +7 mm). In preganglionic, the affected side had decreased HHW, humeral length, and HRC (−0.5 mm, -5%, and -0.7 mm respectively), decreased scapular length (−6%), and tended to have decreased HHT (p=0.065, -0.4 mm). Disarticulation exhibited a smaller GIA (−32 degrees), flatter GRC (+17 mm), decreased HHT and humeral length (−0.4 mm and -31% respectively), and tended to exhibit smaller HRC (p=0.050, -0.3 mm) in the affected side compared to the unaffected.

**Table 3.**
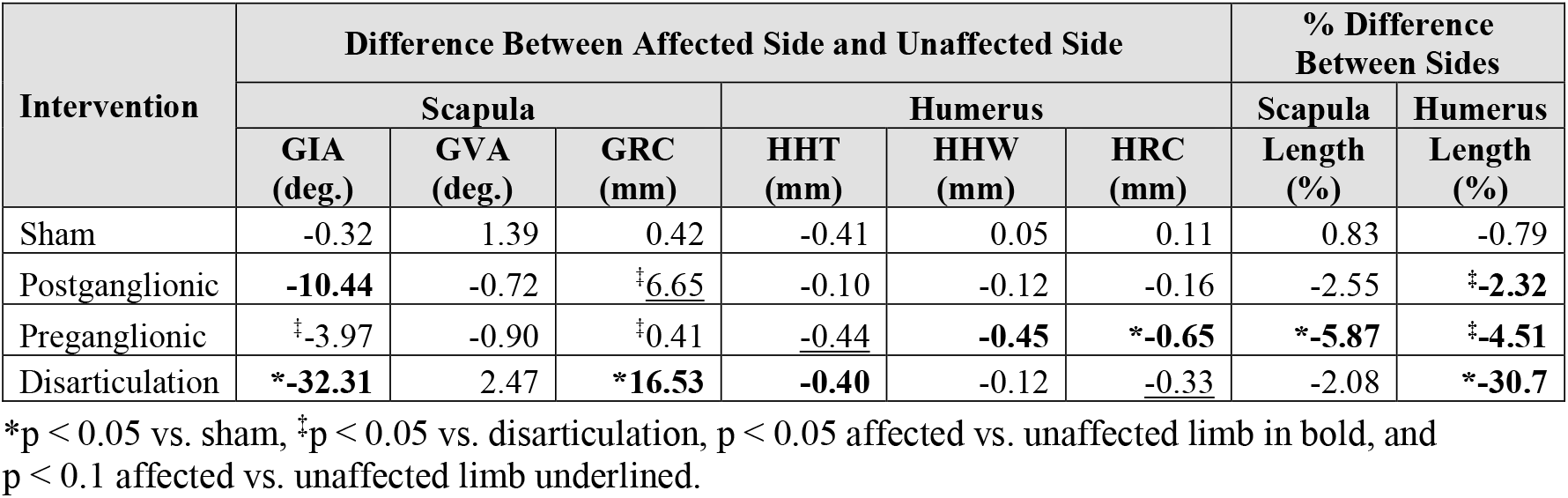
Differences between affected and unaffected limbs in macrostructural metrics in the scapula and humerus. *p < 0.05 vs. sham, ^‡^p < 0.05 vs. disarticulation, p < 0.05 affected vs. unaffected limb in bold, and p < 0.1 affected vs. unaffected limb underlined.

Few metrics were significantly different from sham in the intervention groups (Table 3). Preganglionic showed lower HRC (−0.5 mm) and shorter scapulae (−7%) and disarticulation showed flatter GRC (+16 mm), decreased GIA (+32 degrees), and decreased humeral length (−30%) when compared to sham. Additionally, disarticulation exhibited flatter GRC and shorter humeri than postganglionic (+10 mm and -41% respectively) and preganglionic groups (+16 mm and -38% respectively) and lower GIA compared to preganglionic (−28 degrees).

### 3.3 Multiple Regression Analysis

In general, disarticulation exhibited a greater amount and stronger correlations between the macrostructural and microstructural metrics (Table 4) than the other groups. Between postganglionic and preganglionic groups, postganglionic injury resulted in more correlations between scapular microstructure and scapular morphology that were stronger compared to correlations between humeral microstructure and humeral morphology (Table 5). However, the reverse was true for preganglionic injury. The preganglionic group resulted in more correlations between humeral microstructure and humeral morphology that were stronger compared to correlations between scapular microstructure and scapular morphology (Table 6).

**Table 4.** Top three microstructural predictors of macrostructural morphology and their coefficient of determination in the postganglionic group.

| Morphology |  | Significant Predictors (in order of explanatory power) |  |  |
| --- | --- | --- | --- | --- |
|  |  | 1 | 2 | 3 |
| Glenoid Fossa | Version | -- | -- | -- |
|  | Inclination | Tb.Th<br>(R<sup>2</sup> = 0.337) | +BMD<br>(R<sup>2</sup> = 0.412) | +Conn.D<br>(R<sup>2</sup> = 0.636) |
|  | Curvature | DA<br>(R<sup>2</sup> = 0.278) | +BV/TV<br>(R<sup>2</sup> = 0.452) | -- |
| Scapular Length |  | Tb.Th<br>(R<sup>2</sup> = 0.267) | +DA<br>(R<sup>2</sup> = 0.350) | -- |
| Humeral Head | Thickness | Tb.Th<br>(R<sup>2</sup> = 0.207) | +DA<br>(R<sup>2</sup> = 0.489) | -- |
|  | Width | BMD<br>(R<sup>2</sup> = 0.249) | -- | -- |
|  | Curvature | Conn.D<br>(R<sup>2</sup> = 0.200) | -- | -- |
| Humeral Length |  | -- | -- | -- |

**Table 5.** Top three microstructural predictors of macrostructural morphology and their coefficient of determination in the preganglionic group.

| Morphology |  | Significant Predictors (in order of explanatory power) |  |  |
| --- | --- | --- | --- | --- |
|  |  | 1 | 2 | 3 |
| Glenoid Fossa<sup>a</sup> | Version | -- | -- | -- |
| | Inclination | DA<br>($R^2 = 0.292$) | -- | -- |
| | Curvature | Tb.Th<br>($R^2 = 0.435$) | DA<br>($R^2 = 0.559$) | -- |
| Scapular Length |  | -- | -- | -- |
| Humeral Head<sup>a</sup> | Thickness | Tb.Th<br>($R^2 = 0.500$) | DA<br>($R^2 = 0.616$) | Tb.Sp<br>($R^2 = 0.707$) |
| | Width | BMD<br>($R^2 = 0.471$) | -- | -- |
| | Curvature | BV/TV<br>($R^2 = 0.320$) | Tb.Sp<br>($R^2 = 0.800$) | Tb.N<br>($R^2 = 0.825$) |
| Humeral Length |  | -- | -- | -- |

<sup>a</sup>Gross morphological measurements of the scapula and humerus taken from Dixit *et al.*, 2020.

**Table 6.**
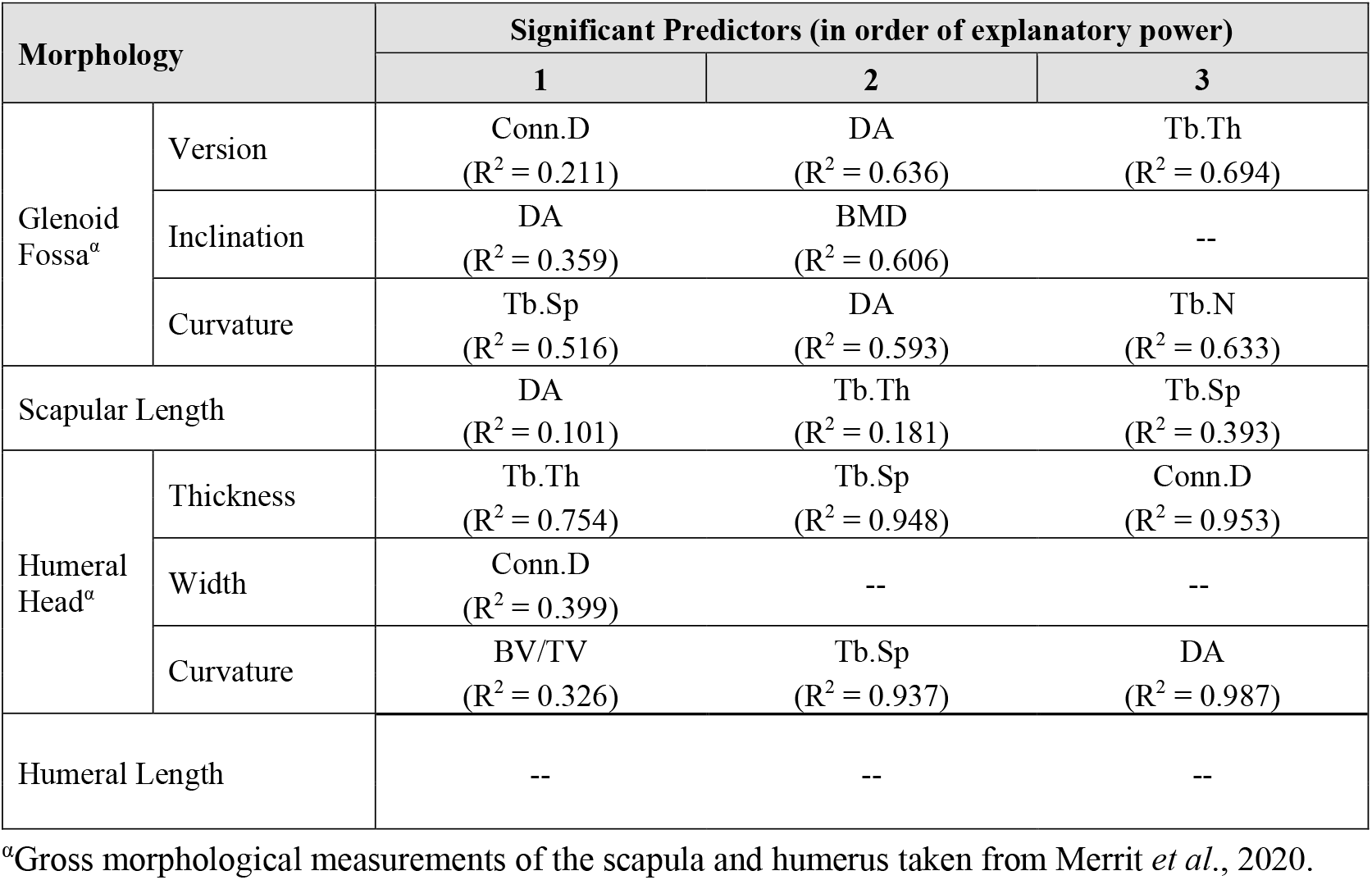
Top three microstructural predictors of macrostructural morphology and their coefficient of determination in the disarticulation group.

| Morphology |  | Significant Predictors (in order of explanatory power) |  |  |
| --- | --- | --- | --- | --- |
|  |  | 1 | 2 | 3 |
| Glenoid Fossa<sup>a</sup> | Version | Conn.D<br>($R^2 = 0.211$) | DA<br>($R^2 = 0.636$) | Tb.Th<br>($R^2 = 0.694$) |
| | Inclination | DA<br>($R^2 = 0.359$) | BMD<br>($R^2 = 0.606$) | -- |
| | Curvature | Tb.Sp<br>($R^2 = 0.516$) | DA<br>($R^2 = 0.593$) | Tb.N<br>($R^2 = 0.633$) |
| Scapular Length | | DA<br>($R^2 = 0.101$) | Tb.Th<br>($R^2 = 0.181$) | Tb.Sp<br>($R^2 = 0.393$) |
| Humeral Head<sup>a</sup> | Thickness | Tb.Th<br>($R^2 = 0.754$) | Tb.Sp<br>($R^2 = 0.948$) | Conn.D<br>($R^2 = 0.953$) |
| | Width | Conn.D<br>($R^2 = 0.399$) | -- | -- |
| | Curvature | BV/TV<br>($R^2 = 0.326$) | Tb.Sp<br>($R^2 = 0.937$) | DA<br>($R^2 = 0.987$) |
| Humeral Length |  | -- | -- | -- |

<sup>a</sup>Gross morphological measurements of the scapula and humerus taken from Merrit *et al.*, 2020.

#### 3.3.1 Scapula

Microstructural metrics from scapular zone 3 are shown to be predictors of scapular morphology, with the strongest correlations in the disarticulation group and the weakest in preganglionic. In postganglionic and preganglionic groups, GVA was not predicted by any of the microstructural metrics. However, in disarticulation, GVA was most predicted by architectural metrics including Conn.D, DA, and Tb.Th (R^2^ = 0.694). In postganglionic, the other morphology metrics were moderately correlated with trabecular quantity and architecture. GIA was best predicted by Tb.Th, BMD, and Conn.D (R^2^ = 0.636). GRC was predicted by DA and BV/TV (R^2^ = 0.452). Scapular length was best predicted by Tb.Th and DA (R^2^ = 0.350). Preganglionic injury showed the weakest and fewest correlations in GIA and GRC with microstructural metrics and showed no correlations with scapular length. GIA was predicted by Tb.Th, BMD, and Conn.D (R^2^ = 0.636). GRC was best predicted by DA and BV/TV (R^2^ = 0.452). The disarticulation group had the strongest and greatest number of correlations in the other scapular macrostructural metrics and was best predicted mainly by trabecular architecture. GIA was predicted by DA and BMD (R^2^ = 0.606). GRC was predicted by Tb.Sp, DA, and Tb.N (R^2^ = 0.633). Scapular length was best predicted by DA, Tb.Th, and Tb.Sp (R^2^ = 0.393). In no groups were morphology metrics correlated with trabecular material properties.

#### 3.3.2 Humerus

Microstructural metrics from the epiphysis are shown to be predictors of humeral head morphology with the strongest correlations in the disarticulation group and the weakest in the postganglionic group. In all injury groups, humeral macrostructure metrics were mainly correlated with trabecular architecture and some trabecular quantity. No morphology metrics were correlated with trabecular material properties and no microstructural metrics were correlated with humeral length. Postganglionic showed weak correlations between epiphyseal microstructure metrics and humeral morphology. HHT was predicted by Tb.Th and DA (R^2^ = 0.489). HHW was predicted by BMD (R^2^ = 0.249) and HRC was predicted by Conn.D (R^2^ = 0.200). Preganglionic showed a greater number of correlations that were stronger between microstructural metrics and humeral morphology. The top three predictors of HHT were Tb.Th, DA, and Tb.Sp (R^2^ = 0.707). As with postganglionic, preganglionic HHW was best predicted by BMD (R^2^ = 0.471). HRC was best predicted by BV/TV, Tb.Sp, and Tb.N (R^2^ = 0.825). The disarticulation group showed even stronger correlations than the other groups. HHT was best predicted by Tb.Th, Tb.Sp and Conn.D (R^2^ = 0.953). HHW differed from the other intervention groups and was best predicted by Conn.D (R^2^ = 0.399). HRC was predicted by BV/TV, Tb.Sp, and DA (R^2^ = 0.987).

## 4. Discussion

In an effort to understand how BPBI affects bone microstructure and its relation to the macrostructural bone deformities, the objective of this study was to characterize the trabecular microstructure within the lateral scapula and proximal humerus and macrostructure morphology of the scapula and humerus following brachial plexus birth injury. Additionally, we compared osseous deficits with regard to injury location with a disuse model to understand how injury location and limb disuse contribute to deformity. Results show substantial deficits following postganglionic BPBI, preganglionic BPBI, and disarticulation in trabecular microstructural throughout the scapula and humerus and some differing macrostructural deficits. More specifically, the lateral scapula had the least robust trabecular bone in zone 3 and the proximal humerus had the least robust trabecular bone in the epiphysis for all intervention groups.

In scapular zone 3, compared to sham, postganglionic had lower trabecular quantity by 16-17% and altered architecture between -2 and 27%, preganglionic had lower trabecular quantity by 32%, material by 3%, and altered architecture between -29 and 42%, and disarticulation had lower trabecular quantity by 19-20% and altered architecture between -30 and 43%. The most profound differences were associated with a preganglionic injury, which had 18-19% lower trabecular quantity and 11% lower Tb.Th compared to postganglionic, 2% less TMD and 19% lower Tb.Th compared to disarticulation, and tended to have 15-17% lower trabecular quantity compared to disarticulation. In the humerus, compared to sham, postganglionic had lower trabecular quantity by 16% and architecture by 6% and preganglionic had lower trabecular quantity by 45-47%, lower TMD by 2%, and altered trabecular architecture between -34 and 65%. Disarticulation had no changes compared to sham. The most profound differences were seen after a preganglionic injury, which had lower quantity by 37% and more altered architecture by -24 to 65% compared to postganglionic, and lower quantity by 38% and more altered architecture by -27 to 45% compared to disarticulation.

These results suggest the overall quality of the bone is diminished in the scapula and humerus following postganglionic injury, preganglionic injury, and disarticulation, but mostly following preganglionic injury. The disarticulation group, representative of limb disuse, aligned more with a postganglionic injury and suggests limb disuse may have a larger effect on postganglionic rather than preganglionic BPBI. Disarticulation also reflected more severe deficits in the scapula versus the humerus and data suggest limb disuse plays a larger role in the scapular deficits compared to the humerus. The results also suggest the overall strength of bone is diminished following BPBI, especially with preganglionic. However, there are likely factors other than disuse affecting strength considered disarticulation did not have as many alterations in bone material property.

Looking at gross morphology, the three intervention groups varied in severity and the bones that were altered. Postganglionic had worse scapular morphology and preganglionic had worse humeral morphology. Disarticulation had worse scapular morphology than even postganglionic but only showed a minor number of detriments in the humerus. Additionally, humeral length was affected in all groups and scapular length only in the preganglionic group. Therefore, it is possible disuse affects humeral length, but factors beyond disuse are causing lack of growth in the preganglionic group.

The correlations between macrostructural and microstructural metrics support the idea of varying degrees of impact from disuse and additional factors beyond limb disuse. Postganglionic injury had weak correlations between metrics in the scapula and weak correlations in the humerus. Preganglionic had moderate correlations between metrics in the humerus and weak correlations in the scapula. However, disarticulation showed the strongest correlations between metrics throughout both the scapula and humerus, implying disuse is affecting both macrostructural and microstructural metrics. In the two neurectomy groups, it is likely the correlations are not as strong because of additional varying factors affecting the macrostructural and microstructural metrics.

Taken together, these results suggest disuse does play a role in trabecular detriments and alterations in gross morphology following BPBI, but is not the only driver. In the scapular glenoid region, disuse had a more profound effect and seems to be a stronger driver for both trabecular deformity and macrostructural alterations. However, the humeral head was not nearly as affected by disuse. Therefore, the effects of nerve injury on the humeral head are not likely to be driven only by disuse, but additional factors such as muscle alterations. The data also show preganglionic trabeculae to be much worse than postganglionic in the humeral head and worse than disarticulation in the scapular neck, suggesting disuse may be a stronger driver in postganglionic BPBI than preganglionic BPBI. Additionally, disuse cannot be the only factor contributing to a loss in strength and overall quality of trabecular bone due to lack of changes in tissue mineral density within the disarticulation group.

Unlike the bone microstructure, the macrostructural osseous deformities have been more fully characterized in both the humerus and the scapula. Many studies agree that glenoid version angle is altered following postganglionic injury [5, 7, 9, 11, 14, 17], but fewer have reported alterations in glenoid inclination angle [5] and glenoid radius of curvature [11, 14] with postganglionic injury. However, the data currently reported on the humeral head is less consistent across studies. Some say the humeral head is flattened [11, 14], some that the head is smaller [5], and some that the head is dislocated or translated [7, 9]. A recent study by our group has data to suggest that the scapular deformities occur more severely in postganglionic injuries while the humeral deformities occur more severely in preganglionic injuries [17]. Our study not only recapitulates the same results as current literature, but also allows for direct comparison between microstructural and macrostructural findings. Overall, our study shows similar results to past data and adds new information about the differences in alterations between injury locations along with how the deformities in bone microstructure and bone macrostructure are associated.

Understanding microstructural and macrostructural osseous deformities in relation to injury location and how it compares to limb disuse is essential to examine potential deformity drivers during postnatal growth. The understanding of how deformities occur will then allow exploration of rehabilitation and prevention of deformities following BPBI. In addition, the differences seen between injury locations can be used to tailor treatments and therapies to a child’s specific needs. Furthermore, the interaction between perinatal peripheral nerve injury and osseous deformity can be applied to a multitude of other diseases and injuries, allowing the scope of the relevance of this data to extend beyond BPBI.

While our study gave insight into injury location, there were some limitations. First, rats are quadrupedal while humans are only quadrupedal before learning to walk. Therefore, the amount of force placed on the glenohumeral joint may differ between the two. However, even with the repetitive loads on the shoulder, the rat model still exhibits similar consequences of nerve injury as in humans [11]. So, although there are slight differences, the model has been confirmed as accurately representing the injury in humans [11]. Second, the timing in which altered trabecular bone can be detected is unknown. Though the age at which the rats in this study were analyzed was before they were skeletally mature, there may be a different age when the trabecular bone is even less robust, whether that be earlier or later than 8 weeks. Future research is needed to examine the timeline in which deformities occur and whether or not they continue to worsen.

## 5. Conclusion

Results suggest postganglionic has worse bone morphology changes, preganglionic has worse bone microstructural changes, and limb disuse is a greater factor for a postganglionic injury. This result implies differing drivers between macrostructural and microstructural alterations, highlighting the need for further investigation behind factors which cause deformity. Prior to this work, it was thought that a postganglionic BPBI was generally worse than a preganglionic BPBI. The knowledge gained here challenges that idea and suggests preganglionic BPBI also causes severe alterations and should be further analyzed. Additionally, the two injury types present different bone deficits and therefore likely require differing treatments. Clinicians need to implement treatments that target the worst bone deformities and deficits likely to occur with the type of nerve injury the patient presents with. However, the specific differences needed in treatment cannot be assessed without first investigating the factors involved in each injury type.

## Acknowledgments

We thank Eric Livingston and Dr. Ted Bateman for micro-CT help. This work was supported by the National Institutes of Health [R21 HD088893] and the National Science Foundation [DGE-1746939].

